# An alternative mode of epithelial polarity in the *Drosophila* midgut

**DOI:** 10.1101/307579

**Authors:** Jia Chen, Aram-Christopher Sayadian, Nick Lowe, Holly E. Lovegrove, Daniel St Johnston

## Abstract

Apical-basal polarity is essential for the formation and function of epithelial tissues, whereas loss of polarity is a hallmark of tumours. Studies in *Drosophila* have identified conserved polarity factors that define the apical (Crumbs, Stardust, Par-6, aPKC), junctional (Baz/Par-3) and basolateral (Scribbled, Discs large, Lgl) domains of epithelial cells *(1, 2)*. Because these conserved factors mark equivalent domains in diverse vertebrate and invertebrate epithelial types, it is generally assumed that this system organises polarity in all epithelia. Here we show that this is not the case, as none of these canonical factors are required for the polarisation of the endodermal epithelium of the *Drosophila* adult midgut. Furthermore, unlike other *Drosophila* epithelia, the midgut forms occluding junctions above adherens junctions, as in vertebrates, and requires the integrin adhesion complex for polarity *(3, 4)*. Thus, *Drosophila* contains two types of epithelia that polarise by different mechanisms. Since knock-outs of canonical polarity factors often have little effect on the polarity of vertebrate epithelia, this diversity of polarity mechanisms is likely to be conserved in other animals *(5-8)*.

---

Current models of epithelial polarity derive from studies on *Drosophila* secretory epithelia, but it is widely believed that this polarity system is universal. Most epithelia form normally in mouse knock-outs of canonical polarity factors, however, with null mutants causing tissue-specific phenotypes that are often unrelated to apical-basal polarity *(3-6).* This has usually been attributed to redundancy between paralogues, but it raises the possibility that some mammalian epithelia may polarise by a different mechanism. This prompted us to ask if all *Drosophila* epithelia polarise in the same way. We therefore examined polarity in the *Drosophila* adult midgut epithelium, which is mainly absorptive rather than secretory and is endodermal in origin, unlike the well-characterised epithelia, which are secretory and arise from the ectoderm or mesoderm *(9)*.

The midgut is a typical epithelium, with an apical brush border marked by actin (Fig. 1C), phospho-Moesin (Fig. 1A) as well as Myosin 1 (Supplementary Fig. 1G) and Myosin 7 (Fig. 1B). However, our analysis led us to rediscover an interesting property of the epithelium: the smooth septate junctions (SJ), which form the occluding barrier to paracellular diffusion, form at the apical side of the lateral domain, above lateral adherens junctions (AJ) *(10, 11)* (Fig. 1D, E). This is the opposite way round to other *Drosophila* epithelia and resembles the junctional arrangement in mammals, where the occluding, tight junctions form above the AJ.

**Figure 1.**
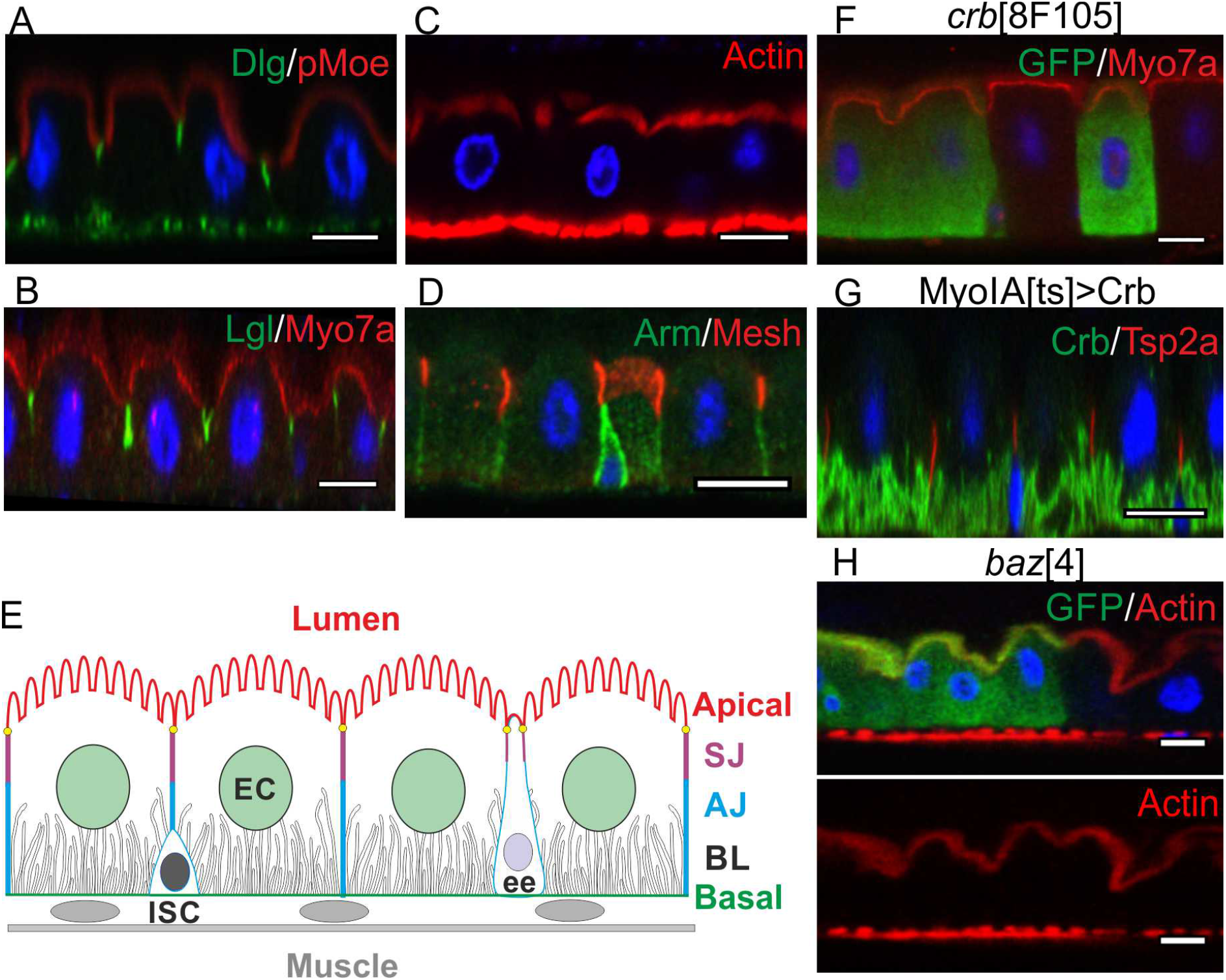
*Drosophila* midgut epithelial cells have a reversed arrangement of junctions to other *Drosophila* epithelia. All panels show apical-basal sections of the *Drosophila* midgut epithelium, with apical on top. **(A)** Phospho-Moesin (red), Dlg (green) and DNA (DAPI; blue) in wild-type. **(B)** Lgl (green) and Myo7a (red). The signal in the enterocyte nuclei in the Myo7a staining is unspecific, since it is present in *ck^13^* (myoVIIa) mutant clones (data not shown). **(C)** Phalloidin-staining of F-actin in the apical brush border. The strong basal signal corresponds to the visceral musculature. **(D)** The adherens junctions, marked by Arm (green) localise basal to the septate junctions, marked by Mesh (red). **(E)** Diagram of the midgut epithelium, which is mainly composed of enterocytes (EC), with a lower frequency of enteroendocrine (ee) cells, both of which turn over in the adult and are replaced by the progeny of basal intestinal stem cell (ISC). The diagram shows the localisation of the septate junctions (SJ) above the adherens junctions (AJ) and the basal labyrinth (BL). **(F)** *crb^8F105^* mutant clones marked by GFP, generated using the Mosaic Analysis with a Repressible Cell Marker (MARCM) technique *(25)*. The apical domain forms normally as revealed by the subcellular localization of Myo7a. **(G)** Conditional mis-expression of UAS-Crb in enterocytes driven by MyoIA-GAL4; tubP-GAL80^*ts*^. Crb localizes to the basal labyrinth and does not perturb apical-basal polarity. Tsp2a is shown in red. **(H)** *baz^4^* mutant cells form a normal apical brush border. Mutant cells are marked by the presence of GFP. Scale bars, 10 μm.

In secretory epithelia, the Crumbs/Stardust complex defines the apical and marginal region and anchors the Par-6/aPKC complex in this domain *(12-14).* Apical Crb and aPKC then exclude Baz/Par-3 to define the apical/lateral boundary by positioning the apical adherens junctions *(15, 16).* This raises the question of whether these factors mark the same positions or the same structures when the junctions are reversed in the midgut. Crumbs is not detectably expressed in the midgut epithelium (Supplementary Fig. 1B) and homozygous mutant clones for null mutations in *crumbs* or *stardust* (*crb^8F-105, 11A22^; sdt^k85^*) give no obvious phenotypes (Fig. 1F and Supplementary Fig. 1E,F). Over-expression of Crumbs expands the apical domain in other *Drosophila* epithelia *(17).* However, ectopically-expressed Crumbs does not affect enterocyte polarity and does not localise apically, concentrating instead in the basal labyrinth (BL), an extensive set of tubular membrane invaginations from the enterocyte basal surface (Fig. 1G). Baz/Par-3 is also not detectable in enterocytes, although it is expressed in the ISCs, where it localises apically (Supplementary Fig. 1A). Consistent with its lack of expression, homozygous clones for *baz* null alleles (*baz^815-8^; baz^4^*) also have no phenotypes (Fig. 1H and Supplementary Fig. 1C,D).

Both aPKC and Par-6 are expressed in the midgut and localise apically, as they do in all other epithelia (Fig. 2A and B). In most polarised cells, the apical localisation of the Par-6/aPKC complex depends on Baz/Par-3, and in epithelia this also requires Crumbs and Stardust (*1*).

**Fig. 2.**
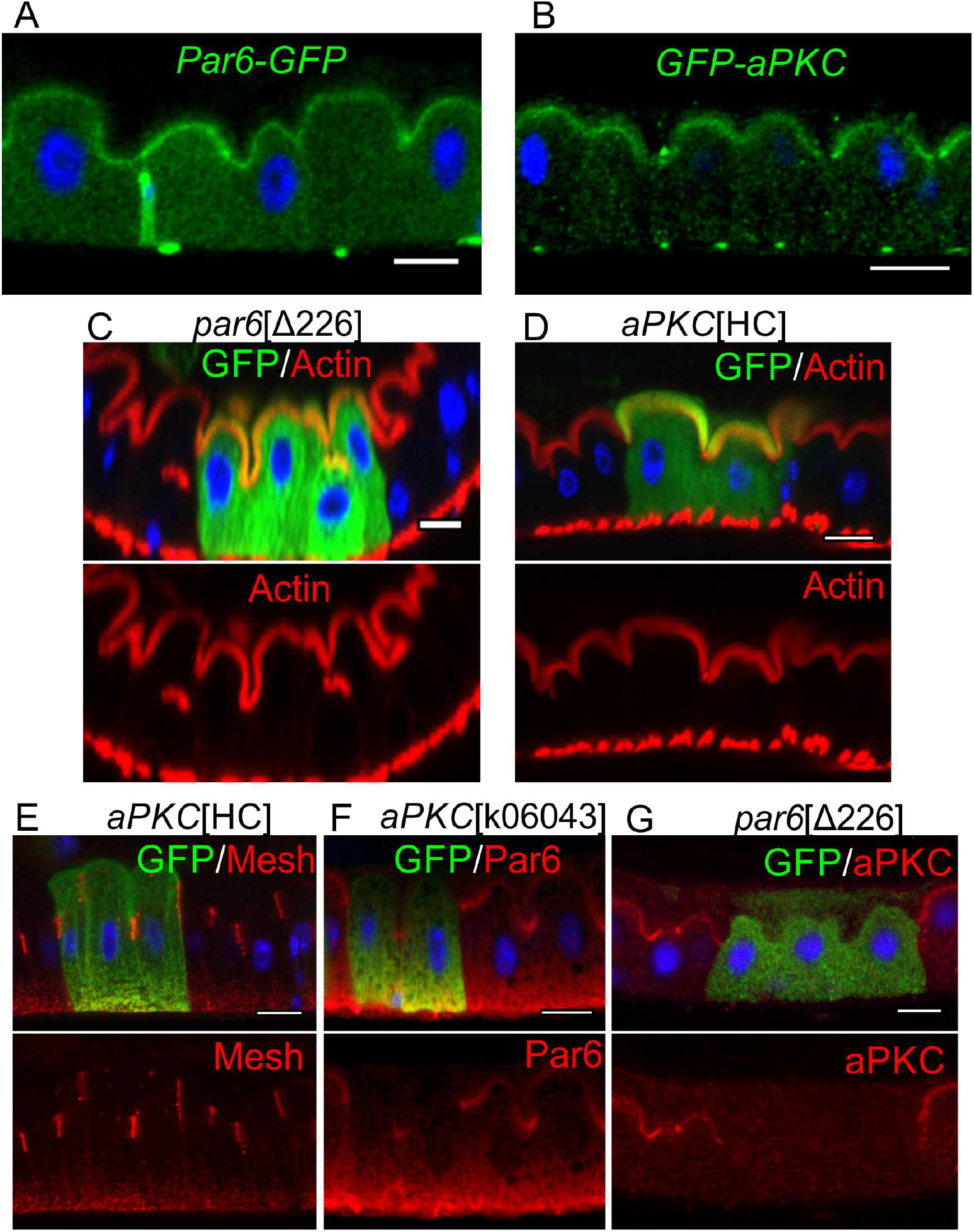
aPKC and Par6 are not required for enterocyte polarity. Subcellular localization of endogenously-tagged Par-6-GFP **(A)** and GFP-aPKC **(B)**, as revealed by anti-GFP staining (green). MARCM clones (marked by GFP) of *par-6*_Δ226_ **(C)** and *aPKC*_HC_ **(D)** show normal apical actin brush borders and SJ localization (**E**; Mesh, red). The apical localization of Par-6 (red) is lost in *aPKC*K06043 MARCM clones **(F)** as is the apical localization of aPKC (red) in *par6*Δ226 MARCM clones **(G)**. Scale bars, 10 μm.

Consistent with our observation that these proteins are absent from enterocytes, *baz* and *crb* mutant clones have no effect on Par-6 localisation, indicating that they must be targeted apically by a distinct mechanism (Supplementary Fig. 2A). Surprisingly, the apical domain forms normally in *par-6^Δ226^* and *aPKC^k06043^* mutant clones, and the morphology of the cells is unaffected (Fig. 2C and data not shown). Although *aPKC^k06043^* is considered a null allele, the corresponding P element insertion does not disrupt the shorter isoforms of aPKC. Thus, it is conceivable that a limited degree of aPKC activity is present in cells homozygous mutant for *aPKC^k06043^*. We therefore used CRISPR to generate a complete null, *aPKC^HC^*, a frameshift mutation that causes a premature stop at amino acid 406. Homozygous *aPKC*^HC^ clones also showed no phenotype, forming normal actin-rich brush borders and septate junctions, confirming that aPKC is dispensable for enterocyte polarity (Fig. 2D,E). Nevertheless, Par-6 is lost from the apical domain in *aPKC^k06004^* clones and aPKC is not apical in *par-6^Δ226^* clones, showing that their localisations are interdependent (Fig. 2F,G). Thus, the Par-6/aPKC complex still marks the apical domain of the midgut epithelium, but is not required for its formation or maintenance.

The lateral polarity factors Scribbled (Scrib), Discs large (Dlg) and Lethal (2) giant larvae (Lgl) are all expressed in the midgut and co-localise with each other and the conserved septate junction component, Coracle (Fig. 1A,B and Supplementary Fig. 2B). Since the septate junctions form at the apical side of the lateral membrane in the midgut, in the position occupied by the adherens junctions in other *Drosophila* epithelia, these proteins mark a conserved structure rather than a conserved position. The lateral epithelial polarity factors are required for septate junction formation in the embryo *(18, 19).* However, the septate junctions form normally in *scrib*, *dlg* and *lgl* mutant clones or when these factors are depleted by RNAi (Fig. 3A,C and Supplementary Fig. 3A). The apical domain is also unaffected in *scrib, dlg* and *lgl* mutant or knock-down cells, in contrast to other epithelia where apical factors are mislocalised to the basolateral domain (Fig. 3E,F and Supplementary Fig.3B,C). Thus, all the canonical epithelial polarity factors are dispensable for the polarisation of the midgut epithelium, even though Par-6, aPKC, Scrib, Dlg and Lgl are expressed and localise to equivalent positions to secretory epithelia.

**Fig. 3.**
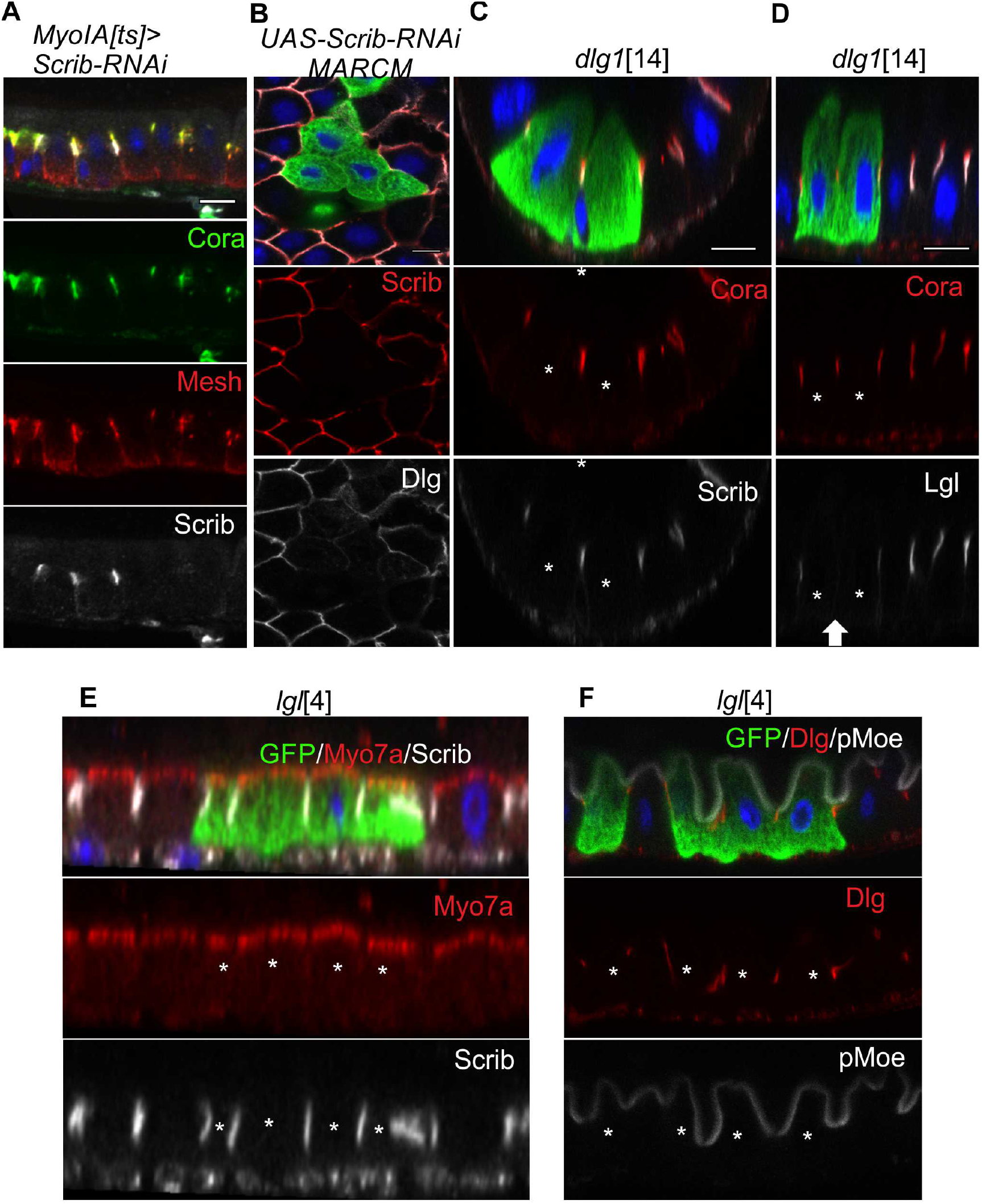
Scrib, Dlg and Lgl localize to septate junctions (SJ) but are not required for SJ formation or apical-basal polarity. **(A)** Mosaic knock-down of Scrib by RNAi in adult enterocytes. The SJ markers Cora (green) and Mesh (red) localize normally in the cells depleted of Scrib (white). **(B)** Dlg (red) does not localize to the SJs in Scrib-RNAi MARCM clones. **(C)** Cora (red) and Scrib (white) localize to the SJs in *dlg1*^14^ MARCM clones. **(D)** Lgl does not localize to the SJ between *dlg1^14^* mutant cells. **(E)** Myo7a (red) and Scrib (white) localize normally to the apical cortex and SJs respectively in *lgl^4^* MARCM clones. **(F)** Dlg (red) and pMoe (white) localize normally in *lgl*^4^ mutant cells. White asterisks mark the mutant cells. The arrow in (d) indicates the SJ between the *dlg1^14^* mutant cells. Scale bars, 10 μm.

The relationships between the lateral factors has been difficult to assess in secretory epithelia because mutants in *scrib, dlg* and *lgl* give rise to round, unpolarised cells without an identifiable lateral domain *(20)*. We took advantage of the normal enterocyte polarisation in these mutants to investigate the interdependence of their recruitment to the septate junctions. Neither Dlg nor Lgl are recruited to septate junctions in cells depleted of Scrib by RNAi (Fig. 3B and Supplementary Fig. 3C). Scrib localises normally in *dlg* mutant clones, whereas Lgl is not localised, and both Scrib and Dlg localise normally to the septate junctions in *lgl* mutant clones (Fig. 3C-F). Thus, there is a simple hierarchical relationship between these factors in the midgut, in which Scrib is required to recruit Dlg, which is needed for Lgl localisation.

The surprising observation that none of the classical epithelial polarity factors are required for the apical-basal polarisation of the midgut raises the question of how polarity is generated and maintained in these cells. Given the similar junctional arrangement to mammalian epithelia, we addressed whether polarity in the midgut depends on integrin-dependent adhesion to the extracellular matrix, as it does in several mammalian epithelia *(7, 21).* Components of the integrin adhesion complex, such as the *α*-integrin Mew and the essential cytoplasmic adaptor proteins, Talin *(Drosophila* rhea) *(22)* and Kindlin *(Drosophila* Fit 1 *(23)*; Fit 2 is not detectable-expressed in the midgut) are highly localised to the basal side of the midgut epithelium (Fig. 4A). The expression of two *α*-integrins and two *β*-integrins in the midgut complicates the genetic analysis of their function, so we focused on the cytoplasmic components of the integrin adhesion complex. Clones of cells homozygous for null alleles of Talin *(rhea^79ã^* and rhea^B128^) detach from the basement membrane and fail to polarise, forming irregularly-shaped cells that do not form septate junctions or an apical domain (Fig. 4B). Most *rhea* mutant cells remain within the epithelial layer, below the septate junctions of the wild-type cells, probably because they do not form septate junctions themselves (Supplementary Fig. 4A).

**Fig. 4.**
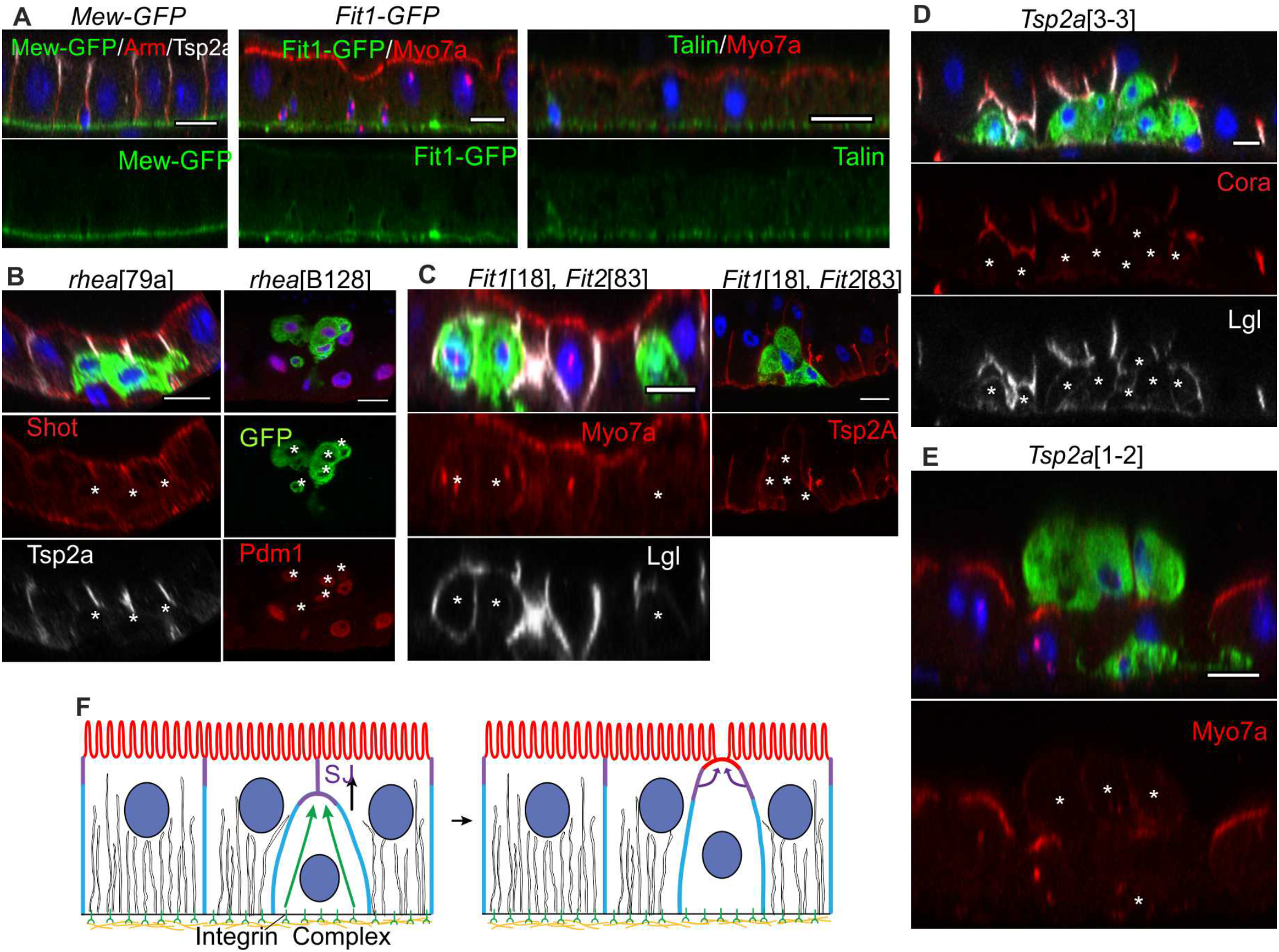
The integrin adhesion complex and septate junction proteins are required for enterocyte polarisation and integration. **(A)** The *α*-integrin Mew and the cytoplasmic adaptor proteins of the integrin adhesion complex, Talin (Rhea) and Fermitin (Fit, also known as Kindlin), localize to the basal surface of the midgut epithelium. Mew-GFP (a protein trap insertion) and Fit-GFP (a genomic fosmid construct) were detected with an anti-GFP antibody (green), whereas the subcellular localization of Talin was revealed with an anti-Talin antibody. **(B)** *rhea*^79a^ mutant cells (marked by GFP) detach from the basement membrane and fail to polarise. Shot (red) is apically enriched in neighbouring wild-type ECs, but is not localized in *rhea* mutant cells, which fail to form SJs marked by Tsp2a (white). Most rhea^B128^ mutant cells express Pdm1 (red), a marker for differentiating ECs. **(C)** *Fit1*^18^ *Fit2*^83^ double mutant clones show a similar phenotype: Myo7a is not enriched apically (red), Lgl (white) spreads around the whole plasma membrane and SJs fail to form as shown by the loss of Tsp2a localization (red). **(D)** *Tsp2a* mutant clones fail to integrate into the epithelium, forming basal clusters that lack SJs, as indicated by the loss of Cora (red) localization and diffuse Lgl staining (white). **(E)** *Tsp2a* mutant cells that face the lumen of the gut do not form an apical domain, as shown by the lack of apical Myo7a (red) enrichment. **(F)** Model of the steps in enterocyte polarisation: adhesion to the basement membrane is required for septate junction formation, which in turn is necessary for the formation of an apical domain. White asterisks * mark the mutant clones. Scale bars, 10 μm.

Despite their inability to polarise and integrate into the epithelium, the mutant cells still appear to differentiate, as they become polyploid and express the marker for differentiating enterocytes, Pdm1 *(24)* (Fig. 4B). Cells mutant for both Fit1 and Fit2 showed a similar phenotype to Talin mutants, whereas mutants or RNAi of other components of the integrin adhesion complex had no obvious effect (Fig. 4C, Supplementary Fig. 4B,C and data not shown).

The intestinal stem cells lie beneath the epithelium and differentiating enterocytes must therefore integrate into the epithelium from the basal side, inserting between the septate junctions of the flanking enterocytes, while maintaining an intact barrier. We therefore examined the effects of mutants in the core septate junction components, Tsp2a and Mesh *(11)*. More than 90% of mutant cells fail to integrate through the septate junctions of the neighbouring wild-type cells, and the clones form cysts on the basal side of the epithelium (Fig. 4D and Supplementary Fig. 4A,D). Wild-type cells start to form an apical domain as they integrate, before they have a free apical surface, as shown by the enrichment of apical components, such as Myo7a (Supplementary Fig. 4E). By contrast, apical markers never localise into a clear apical domain in *Tsp2a* mutant cells, even if the cells are extruded from the apical side of epithelium (Fig. 4E and Supplementary Fig. 4E). Thus, the midgut epithelium polarises in a basal to apical manner, in which adhesion to the ECM is required for the formation of the septate junctions, and the septate junctions are needed for the formation of the apical domain (Fig. 4F).

Our results reveal that the idea that there is a universal system for polarising epithelial cells is incorrect, as the polarity of the intestinal epithelium is fundamentally different from other *Drosophila* epithelia, indicating that flies have two classes of epithelia that polarise by distinct mechanisms. This difference is unlikely to reflect the fact that the midgut is absorptive rather than secretory, as secretory cells in the midgut, such as the enteroendocrine cells and the acid-secreting copper cells, polarise in the same way as the enterocytes (data not shown). The different polarity mechanism of the midgut could be a consequence of its developmental origin, since it arises from the endoderm, whereas all other epithelia are ectodermal or mesodermal, or it may reflect its reversed arrangement of occluding and adherens junctions compared to other epithelia. Alternatively, it may correspond to the direction in which the cells polarise, as enterocytes polarise in a basal to apical direction as they integrate into the epithelium, whereas other *Drosophila* epithelia acquire polarity in an apical to basal direction.

Polarity in the *Drosophila* midgut resembles that of well-characterised mammalian epithelia in its arrangement of junctions, the lack of a requirement for classical polarity factors and its dependence on the integrin adhesion complex, suggesting that it will be a good *in vivo* model for at least some types of vertebrate epithelia. It will therefore be important to determine whether vertebrates also contain two types of epithelia whose polarity is controlled by different factors.

## Materials & Methods

### *Drosophila* genetics

*w*[1118] or *y*[2] flies were used as wild-type unless otherwise specified. Other stocks used in this study were:

**Fluorescently-tagged protein lines:** EGFP-aPKC (this study), Par6-EGFP*(26)*, Mew-YFP*(27)* (Kyoto DGRC # 115524), Baz-EGFP*(28)* (Bloomington # 51572), Myo31DF-YFP*(27)* (Kyoto DGRC # 115611), Crb-EGFP *(29)* (gift from Y. Hong, University of Pittsburgh, USA) Fit1-EGFP (gift from B. Klapholz and N. Brown, Department of Physiology, Development and Neuroscience, University of Cambridge, UK).

**Mutant stocks:** *aPKC[k06043]*(30)*, aPKC[HC]* (this study), *par6[Δ226]*(31)*, dlg1[A]* (Bloomington # 57086), *dlg1[14](32)*, *lgl[4](33)*, *baz[815-8](34)*, *baz[4](35)*, *crb[8F105](36)*, *crb[11A22](36)*, *Tsp2A[1-2]*, *Tsp2A[3-3]*, *Tsp2A[2-9](37)* (gift from M. Furuse, Kobe University, Japan), *mesh[f04955](38)* (Kyoto DGRC # 114660), *Fit1[18]*, *Fit2[83]*, *Fit1-EGFP*, *Fit1[18] Fit2[83](23)*, *rhea[79a](22), rhea[B28]*, *rhea[B128](23)*, *ilk[54](39)* (gifts from B. Klapholz and N. Brown, Department of Physiology, Development and Neuroscience, University of Cambridge, UK).

**UAS responder lines:** UAS-Scrib-RNAi (TRiP; Bloomington # 35748), UAS-Crb (Bloomington # 5544).

The following stocks were used to generate (positively labeled) MARCM clones:

MARCM FRTG13: y w, UAS-mCD8::GFP, Act5C-GAL4, hsFLP[1]; FRTG13 tubP-GAL80 MARCM FRT82B: y w, UAS-mCD8::GFP, Act5C-GAL4, hsFLP[1];; FRT82B tubP-GAL80 MARCM FRT19A: w, hsFLP, tubP-GAL80, FRT19A;; tubP-GAL4, UAS-mCD8::GFP / TM3, Sb

MARCM FRT2A: hsFLP[1]; tubP-GAL4, UAS-mCD8::GFP / CyO, GFP; FRT2AtubP-GAL80 (gift from B. Klapholz and N. Brown)

MARCM FRT40A: y w, UAS-mCD8::GFP, Act5C-GAL4, hsFLP[1]; FRT40A tubP-GAL80.

Negatively marked clones on the X chromosome were generated using the following stock: y w His2Av::GFP hsFLP[12] FRT19A / FM7a (Bloomington # 32045).

Clones mutant for *mesh* were generated using the following stock: esg-GAL4, UAS-FLP, tubP-GAL80[ts] / CyO; FRT82B nlsGFP (referred to as esg>Flp in Supplementary Fig. 4d; gift from G. Kolahgar, Department of Physiology, Development and Neuroscience, University of Cambridge, UK).

UAS-RNAi constructs and UAS-Crb were expressed in the adult midgut epithelium using the following driver line: y w; MyoIA-GAL4, tubP-GAL80[ts] (referred to as Myo1[ts] in Fig. 1g, 3a,b, and Supplementary Fig. 3c; gift from G. Kolahgar).

### Stock Maintenance

Standard procedures were used for *Drosophila* husbandry and experiments. Flies were reared on standard fly food supplemented with live yeast at 25 °C. For the conditional expression of UAS responder constructs (e.g. RNAi), parental flies were crossed at 18 °C and the resulting offspring reared at the same temperature until eclosion. Adult offspring were collected for 3 days and then transferred to 29 °C to inactivate the temperature-sensitive GAL80 protein. To generate MARCM or GFP-negative clones, flies were crossed at 25 °C and the resulting offspring subjected to heat-shocks either as larvae (from L2 until eclosion) or as adults (5 - 9 days after eclosion). Heat shocks were performed at 37 °C for 1 h (twice daily). Flies were transferred to fresh food vials every 2 - 3 days and kept at 25 °C for at least 9 days after the last heat shock to ensure that all wild-type gene products from the heterozygous progenitor cells had turned over. For this study, all (midgut) samples were obtained from adult female flies.

### Formaldehyde fixation

Samples were dissected in PBS and fixed with 8% formaldehyde (in PBS containing 0.1% Triton X-100) for 10 min at room temperature. Following several washes with PBS supplemented with 0.1% Triton X-100 (washing buffer), samples were incubated in PBS containing 3% normal goat serum (NGS, Stratech Scientific Ltd, Cat. # 005-000-121; concentration of stock solution: 10 mg / ml) and 0.1% Triton X-100 (blocking buffer) for 30 min. This fixation method was only used for samples in which F-actin was stained with fluorescently-labelled phalloidin, as phalloidin staining is incompatible with heat fixation.

### Heat fixation

The heat fixation protocol is based on a heat-methanol fixation method used for *Drosophila* embryos*(40)*. Samples were dissected in PBS, transferred to a wire mesh basket, and fixed in hot 1X TSS buffer (0.3% Triton X-100, 0.4 g / L NaCl; 95 °C) for 3 sec, before being transferred to ice-cold 1X TSS buffer and chilled for at least 1 min. Subsequently, samples were transferred to washing buffer and processed for immunofluorescence stainings

### Immunofluorescence

After blocking, samples were incubated with the appropriate primary antibody / antibodies diluted in blocking buffer at 4 °C overnight. Following several washes, samples were incubated with the appropriate secondary antibody / antibodies either at room temperature for 2 h or at 4 °C overnight. Samples were then washed several times in washing buffer and mounted in Vectashield containing DAPI (Vector Laboratories) on borosilicate glass slides (No. 1.5, VWR International). All antibodies used in this study were tested for specificity using clonal analysis (MARCM) or RNAi.

#### Primary antibodies

Mouse monoclonal antibodies: anti-Dlg (4F3), anti-Cora (c615.16), anti-αSpec (3A9), anti-Arm (N2 7A1), anti-Talin (A22A, E16B), anti-Pros (MR1A), anti-Crb (Cq4), anti-Nrv (Nrv5F7), anti-Mys (CF.6G11). All monoclonal antibodies were obtained from the Developmental Studies Hybridoma Bank and used at 1:100 dilution.

Rabbit polyclonal antibodies: anti-pEzrin (NEB Cat. # 3726S, 1:200 dilution); anti-Lgl (Santa Cruz Biotechnoloy Inc., d-300, Cat. # SC98260, 1:200 dilution); anti-aPKC (Santa Cruz Biotechnoloy Inc., Cat # SC216, 1:100 dilution); anti-*β*_H_Spec*(47)* (gift from C. Thomas, The Pennsylvania State University, USA, 1:1000 dilution); anti-Bazooka*(30)* (gift from A. Wodarz, University of Cologne, Germany, 1:200 dilution); anti-Par6*(42)* (gift from D. J. Montell, UCSB, USA, 1:500 dilution); anti-Mesh*(38)* and anti-Tsp2A*(37)* (gift from M.

Furuse, 1:1000 dilution); anti-Scrib*(43)* (gift from C. Q. Doe, University of Oregon, USA, 1:1000 dilution); anti-Pdm1*(44)* (gift from F. J. Diaz-Benjumea, Centre for Molecular Biology "Severo Ochoa" (CBMSO), Spain, 1:1000 dilution); anti-Cno*(45)* (gift from M. Peifer, UNC, USA, 1:1000 dilution).

Other antibodies used: chicken anti-GFP (Abcam, Cat. # ab13970, 1:1000 dilution); guinea pig anti-Myo7a*(46)* (gift from D. Godt, University of Toronto, Canada, 1:1000 dilution); guinea pig anti-Shot*(47)* (1:1000 dilution); rat anti-Mesh*(38)* (gift from M. Furuse, 1:1000 dilution).

#### Secondary Antibodies

Alexa Fluor secondary antibodies (Invitrogen) were used at a dilution of 1:1000.

Alexa Fluor 488 goat anti-mouse (# A11029), Alexa Fluor 488 goat anti-rabbit (# A11034), Alexa Fluor 488 goat anti-guinea pig (# A11073), Alexa Fluor 488 goat anti-chicken IgY (# A11039), Alexa Fluor 555 goat anti-rat (# A21434), Alexa Fluor 555 goat anti-mouse (# A21422), Alexa Fluor 555 goat anti-rabbit (# A21428), Alexa Fluor 568 goat anti-guinea pig (# A11075), Alexa Fluor 647 goat anti-mouse (# A21236), Alexa Fluor 647 goat anti-rabbit (# A21245), Alexa Fluor 647 goat anti-rat (# A21247). Only cross-adsorbed secondary antibodies were used in this study to eliminate the risk of cross-reactivity.

F-Actin was stained with phalloidin conjugated to Rhodamine (Invitrogen, Cat. # R415, 1:500 dilution).

### Imaging

Images were collected on an Olympus IX81 (40X 1.35 NA Oil UPlanSApo, 60X 1.35 NA Oil UPlanSApo) using the Olympus FluoView software Version 3.1 and processed with Fiji (ImageJ).

### Generation of endogenous EGFP-aPKC

Endogenously tagged aPKC with EGFP fused to the N-terminus was generated by CRISPR mediated homologous recombination. *In vitro* synthesised gRNA *(48)* to a CRISPR target approximately 60 nucleotides downstream from the *aPKC* start codon (target sequence GAATAGCGCCAGTATGAACATGG) and a plasmid donor containing the ORF of EGFP as well as appropriate homology arms (1.5 kb upstream and downstream) were co-injected into nos-Cas9 expressing embryos (Bloomington # 54591; also known as CFD2)*(49)*. Single F0 flies were mated to *y w* flies and allowed to produce larvae before the parent was retrieved for PCR analysis. Progeny from F0 flies in which a recombination event occured (as indicated by PCR) were further crossed and analysed to confirm integration. Several independent EGFP-aPKC lines were isolated. Recombinants carry the EGFP coding sequence inserted immediately downstream of the endogenous start codon and a short linker (amino acid sequence: Gly Ser Gly Ser) between the coding sequence for EGFP and the coding sequence for aPKC. Homozygous flies are viable and healthy.

### Generation of *aPKC[HC]*

We used the CRISPR/Cas9 method*(48)* to generate a null allele of *aPKC.* sgRNA was *in vitro* transcribed from a DNA template created by PCR from two partially complementary primers: forward primer: 5’-

GAAATTAATACGACTCACTATAggattacggcatgtgtaaggGTTTTAGAGCTAGAAATAGC-3’; reverse primer: 5’-

AAAAGCACCGACTCGGTGCCACTTTTTCAAGTTGATAACGGACTAGCCTTATTTT AACTTGCTATTTCTAGCTCTAAAAC-3’. The sgRNA and Cas9 mRNA were co-injected into nos-Cas9 embryos. Putative *aPKC* mutants in the progeny of the injected embryos were recovered, balanced, and sequenced. The *aPKC[HC]* allele contains a small deletion around the CRISPR site, resulting in one missense mutation and a frameshift that leads to stop codon at amino acid 406 in the middle of the kinase domain, which is shared by all isoforms (see below). The *aPKC[HC]* allele was subsequently recombined onto FRTG13 to generate MARCM clones. No aPKC protein was detectable by antibody staining in both midgut and follicle cell clones, and follicle cells homozygous mutant for *aPKC[HC]* display a phenotype that is indistinguishable from that observed in follicle cells homozygous mutant for the *aPKC[K06403]* allele (data not shown).

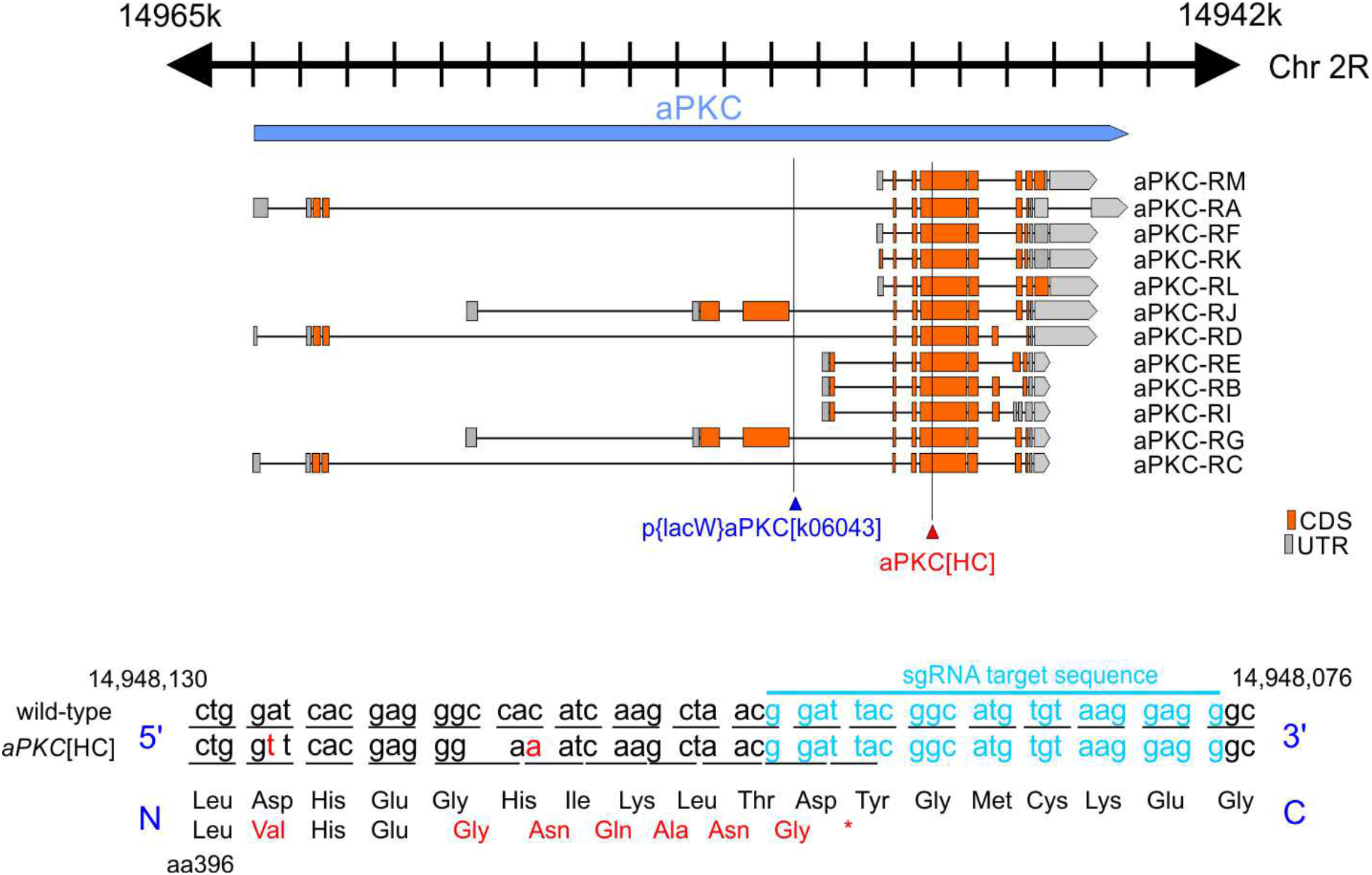

### Statistics

The proportions of *rhea, Fit* and *Tsp2a* mutant cells inside the epithelial layer were calculated as follows: images were taken of different regions of several midguts containing MARCM clones stained with an apical marker. Cells that were above the neighbouring cells or had a clear apical domain were counted as “cells NOT inside the layer”, whereas cells without a detectable free apical surface were counted as “cells inside the layer”. Data were analysed with Graphpad Prism software. The graph in Supplementary Fig. 4a shows the average % of cells inside the layer ± SD.

### Reproducibility of experiments

All experiments were repeated multiple times with independent crosses.

Baz-EGFP (4 independent experiments), EGFP-aPKC (9 independent experiments), Mew-YFP (5 independent experiments), Par6-EGFP (4 independent experiments), Crb-EGFP (5 independent experiments), Fit1-EGFP (4 independent experiments), Myo31DF-YFP (4 independent experiments), MyoIA[ts]>UAS-Scrib-RNAi (11 independent experiments), MyoIA[ts]>UAS-Crb (6 independent experiments).

The phenotypes of homozygous mutant clones were analysed in multiple guts from independent experiments as follows: *baz[4]* (13 independent experiments; 3078 mutant cells analysed), *baz[815-8]* (4 independent experiments; 734 cells analysed), *aPKC[k06043]* (8 independent experiments; 3681 mutant cells analysed), *aPKC[HC]* (9 independent experiments; 15984 mutant cells analysed), *par6[ħ>226]* (4 independent experiments; 2558 mutant cells analysed), *crb[11A22]* (4 independent experiments; 1478 mutant cells analysed), *crb[8F105]* (5 independent experiments; 3288 mutant cells analysed), *lgl[4]* (6 independent experiments; 6790 mutant cells analysed), *dlg1[14]* (5 independent experiments; 3092 mutant cells analysed), *rhea[79a]*, rhea[B28], *rhea[B128]* (4 independent experiments for each genotype; 175 mutant cells analysed in total), *Fit1[18], Fit2[83]* (7 independent experiments; 608 double mutant cells analysed), *Fit1[18]* (4 independent experiments; 854 mutant cells analysed), *ilk[54]* (5 independent experiments; 65 mutant cells analysed), *Tsp2a[1-2] Tsp2a[2-9]* and *Tsp2a[3-3]* (7, 4 and 6 independent experiments respectively; a total of 1205 *Tsp2a* mutant cells were analysed), *mesh[f04955]* (5 independent experiments; 643 mutant cells analysed).

## Acknowledgments

We are grateful to C. Mendes for discussions and help with experiments; G. Kolahgar, Y. Izumi and M. Furuse, B. Klapholz and N. Brown for fly lines and reagents; C. Thomas, A. Wodarz, D. J. Montell, C. Q. Doe, F. J. Diaz-Benjumea, M. Peifer, and D. Godt for antibodies; members of the St Johnston laboratory for discussions and comments on the manuscript.

## Funding

This work was supported by a Wellcome Trust Principal Fellowship to D.St J. (080007, 207496) and by core support from the Wellcome Trust (092096, 203144) and Cancer Research UK (A14492, A24823). J.C. was supported by Royal Society K.C. Wong Postdoctoral Fellowship. A.C.S. was supported by a Wellcome Trust 4 year PhD studentship (083348).

## Author contributions

J.C., A.C.S. and D.St J. designed the experiments; J.C. and A.C.S. conducted all experiments and analyses; J.C., A.C.S. and D.St J. wrote the manuscript; N.L. generated the EGFP-aPKC strain; H.L. generated the *aPKC[HC]* mutation.

---

**Supplementary Fig1.**
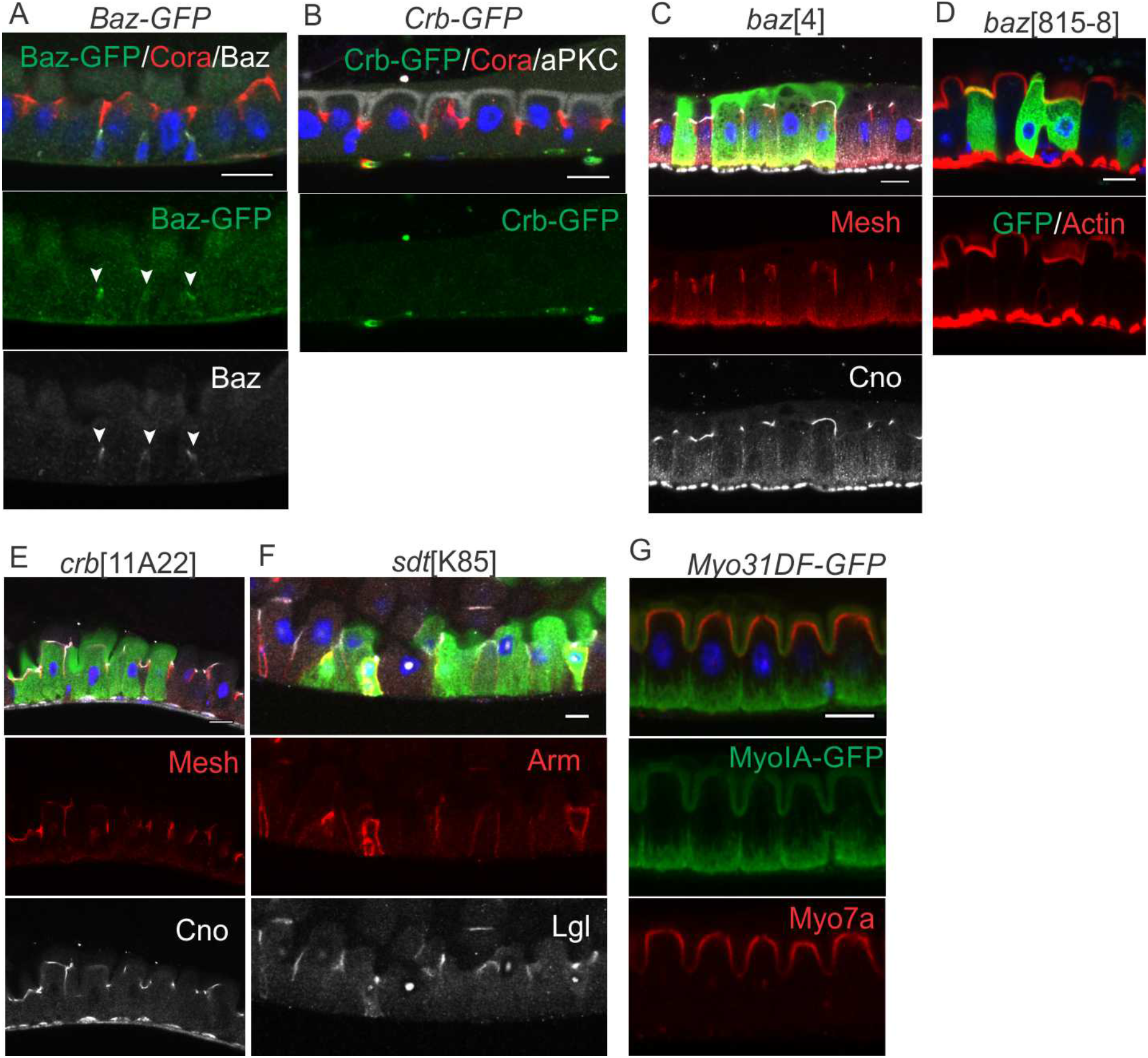
Baz and Crb are not detectably expressed in midgut enterocytes and are not required for apical-basal polarity in the midgut epithelium. **(A)** Baz protein trap line stained for GFP (green), Cora (red) and Baz (white). The arrowheads mark Baz in the ISCs, where it localises apically. **(B)** Crb protein trap line stained for GFP (green), Cora (red) and aPKC (white). **(C)** *baz*^4^ MARCM clones (marked by GFP) show normal Mesh (red) and Cno (white) localization. **(D)** *baz^8l5-8s^* mutant cells (marked by GFP) form a normal apical brush border as revealed by phalloidin staining of F-actin (red). **(E)** *crb^11A22^* MARCM clones show normal Mesh (red) and Cno (white) localisation. **(F)** *sdt^K85^* MARCM clones show normal Arm (red) and Lgl (white) localisation. (G) Myo31DF/MyoIA protein trap line stained for GFP (green) and Myo7a (red). Scale bars, 10 μm.

**Supplementary Figure 2.**
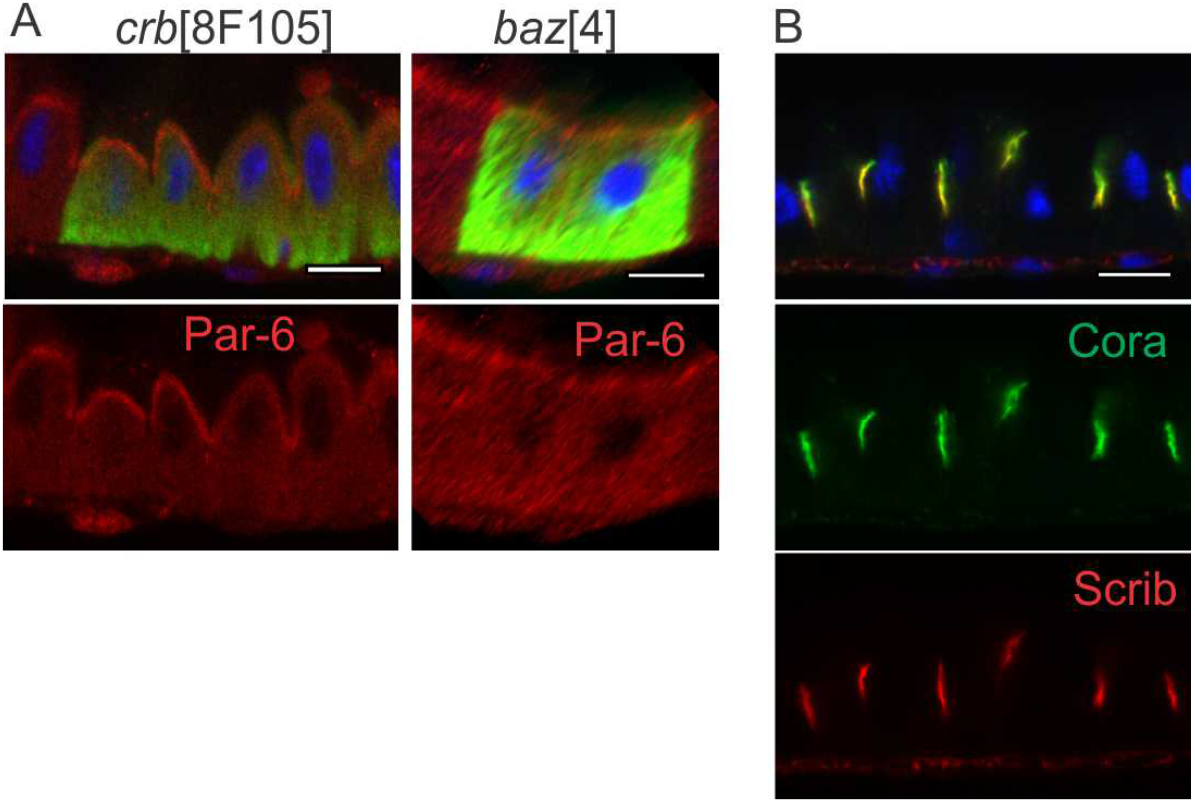
Crumbs and Bazooka are not required to recruit Par-6 apically. **(A)** Par-6 localises normally to the apical surface of *crb^Fl05^* and *baz^4^* mutant cells (marked by GFP). Scrib (red) localises to the enterocyte SJs, marked by Cora (green). Scale bar, 10 μm.

**Supplementary Figure 3.**
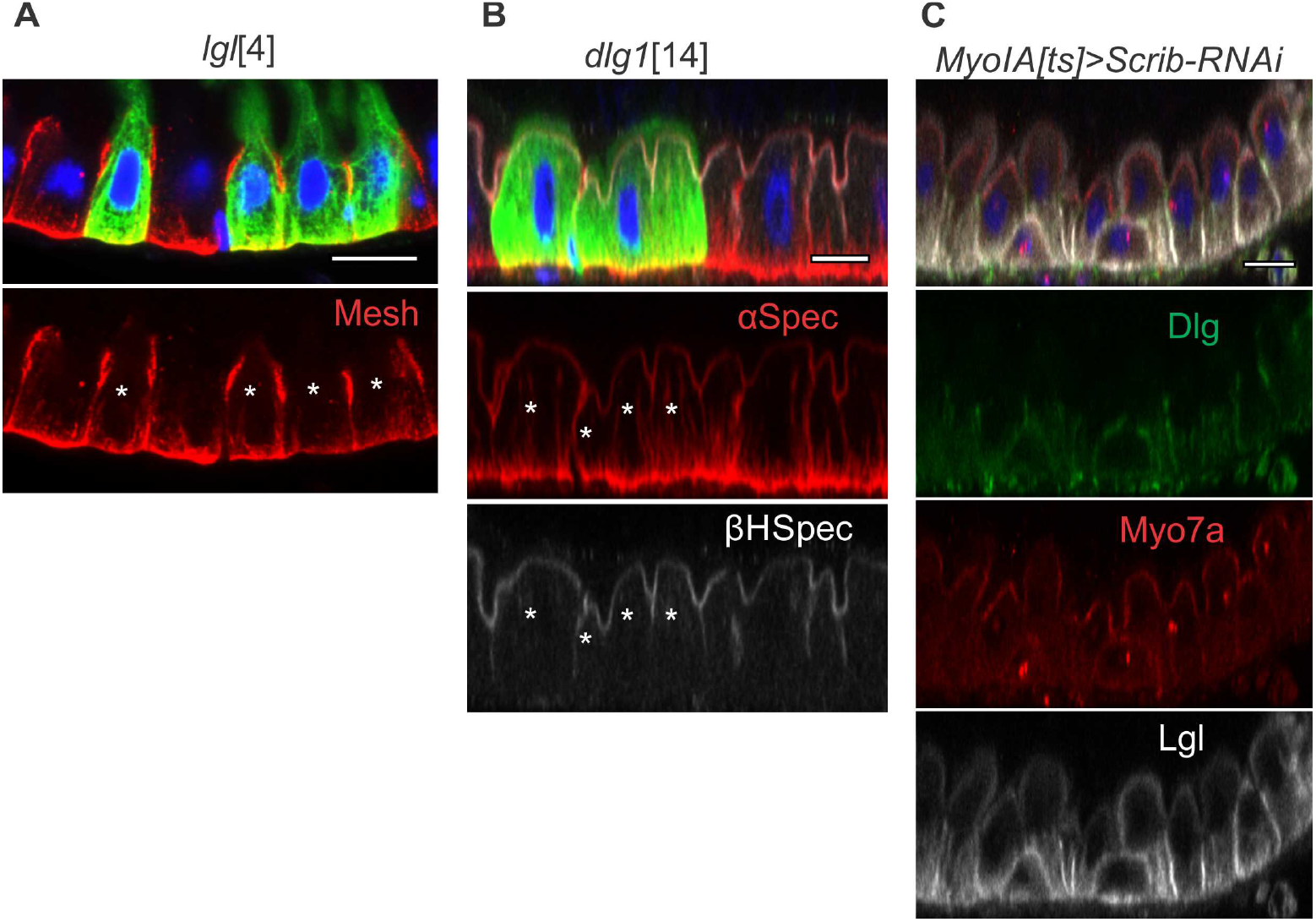
Loss of Scrib or Dlg has no effect on apical domain formation in enterocytes. **(A)** Mesh (red) localises normally to the SJs of *lgf* mutant cells (marked by GFP). (B) *dlg1^u^* MARCM clones show normal apical localization of a-Spectrin (red) and *β*_H_-Spectrin (white). (C) RNAi knock-down of Scrib in adult midgut enterocytes has no effect on the subcellular localization of Myo7a (red), but disrupts Dlg (green) and Lgl (white) localisation to the SJs. Scale bar, 10 μm.

**Supplementary Fig 4.**
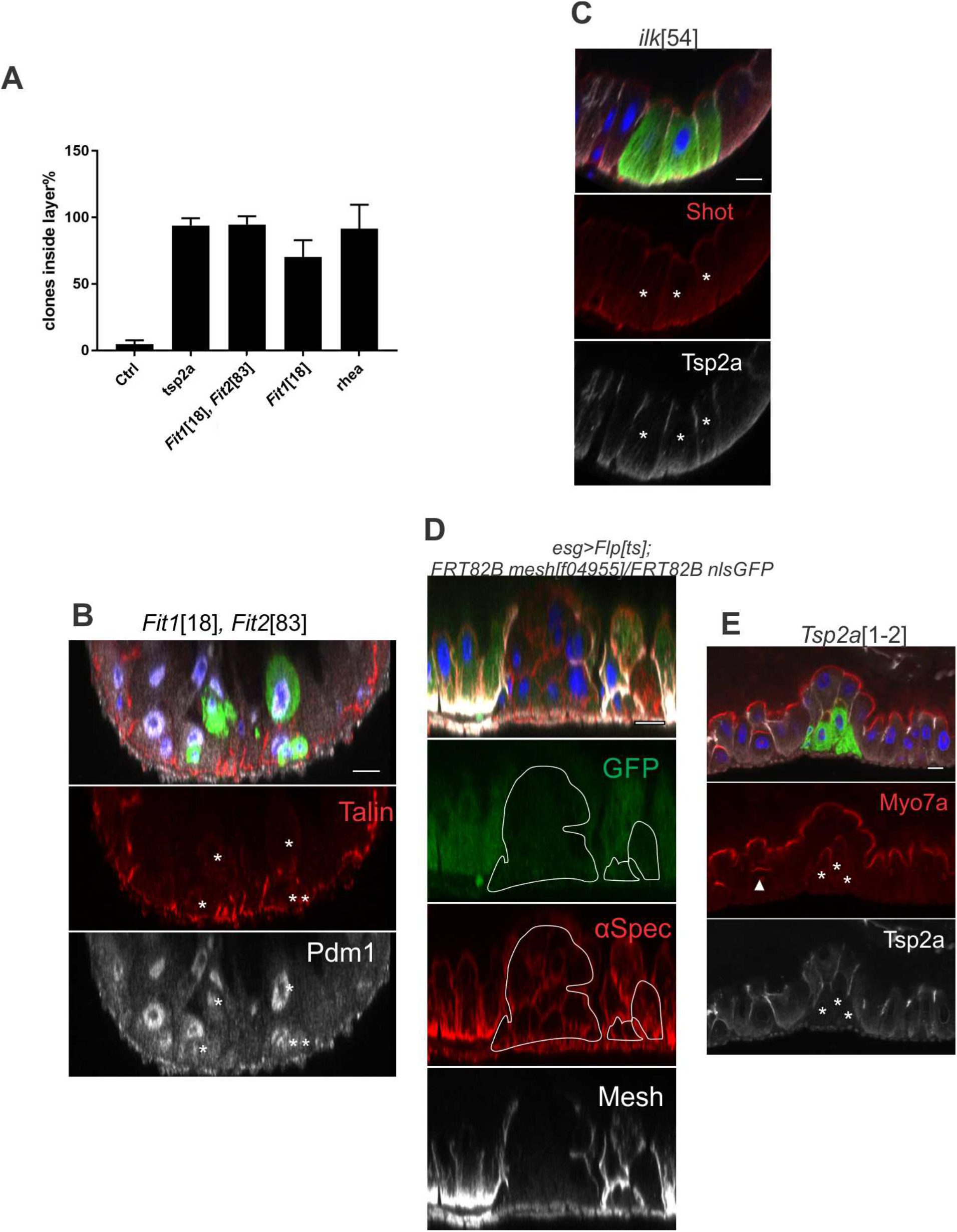
Talin, Kindlin and SJ components are required for enterocyte polarity. **(A)** Most *rhea, Fit1, Fit1 Fit2*, and *Tsp2a* mutant cells remain inside the epithelia layer. The graph is based on the analysis of 299 cells in wild-type MARCM clones (14 images), 1205 *Tsp2a* mutant cells (27 images from *Tsp2a^1-2^*, *Tsp2a^3-3^* and *Tsp2a^2-9^* clones), 608 *Fit1*^18^ *Fit2*^83^ (*Fit^D^*) double mutant cells (24 images), 854 *Fit1*^18^ mutant cells (25 images) and 175 *rhea* mutant cells (22 images from *rhea^B28^*, *rhea^19^* and *rhea* ^B128^). **(B)** *Fit1*^18^ *Fit2^88^* double mutant cells differentiate as ECs, as revealed by the expression of Pdm1 (white). **(C)** *ilk^54^* mutant cells have normal polarity, as revealed by the apical enrichment of Shot (red) and the localisation of Tsp2a (white) to the SJs. **(D)** A *mesh^m955^* mutant clone (marked by the loss of GFP) stained for anti-a-Spectrin (red) and Mesh (white). **(E)** A *Tsp2a^1-2^* MARCM clone showing weak and diffuse Myo7a (red) localisation compared to the strong apical enrichment seen in neighbouring wild-type cells that are inserting into the epithelium (white arrow head). White asterisks * and lines mark the mutant clones. Scale bars, 10 μm.

